## Supporting information material (description in the main manuscript) for "Objects guide human gaze behavior in dynamic real-world scenes": S1_tab.pdf

**Table S1.** List of the 23 videos of the VidCom dataset used in this study. The column “*Subjects”* shows the number of human observers, each contributing to ground truth scanpaths in the human data, and “*Split*” indicates if the video was used in the training or test set.

| **Name** | **Subjects** | **Split** |
|---|---|---|
| dance01 | 12 | train |
| dance02 | 14 | train |
| field03 | 12 | test |
| foutain02 | 14 | test |
| garden04 | 12 | test |
| garden06 | 14 | train |
| garden07 | 13 | train |
| garden09 | 12 | test |
| park01 | 12 | train |
| park06 | 12 | train |
| park09 | 11 | test |
| road02 | 11 | train |
| road04 | 11 | test |
| road05 | 11 | train |
| robarm01 | 12 | test |
| room01 | 10 | train |
| room02 | 12 | test |
| room03 | 10 | test |
| tommy02 | 13 | test |
| uscdog01 | 12 | test |
| walkway01 | 13 | test |
| walkway02 | 13 | test |
| walkway03 | 13 | train |
