## Supplementary figures and images for "Objects guide human gaze behavior in dynamic real-world scenes"

### S1_fig.tif

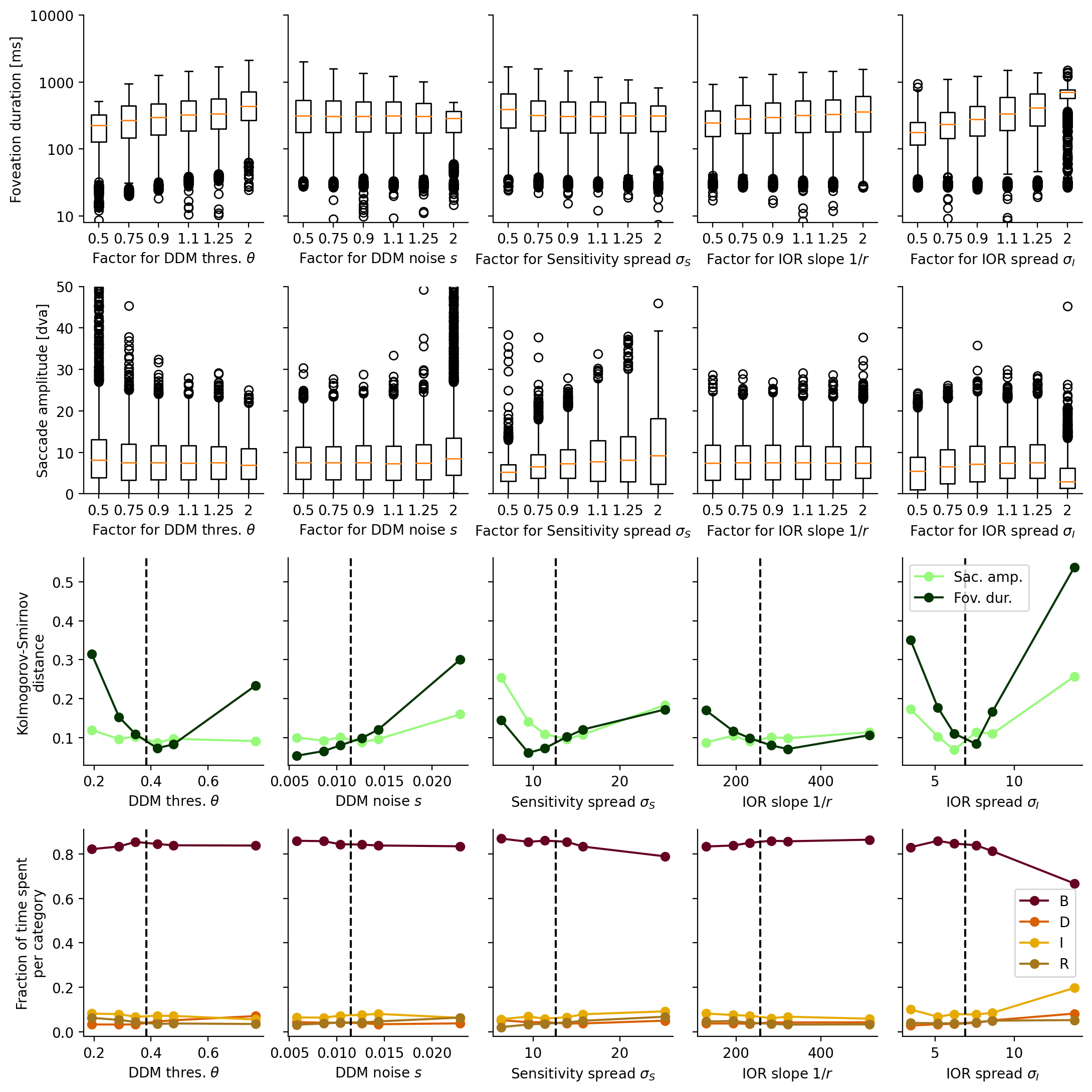

### S1_file.gif

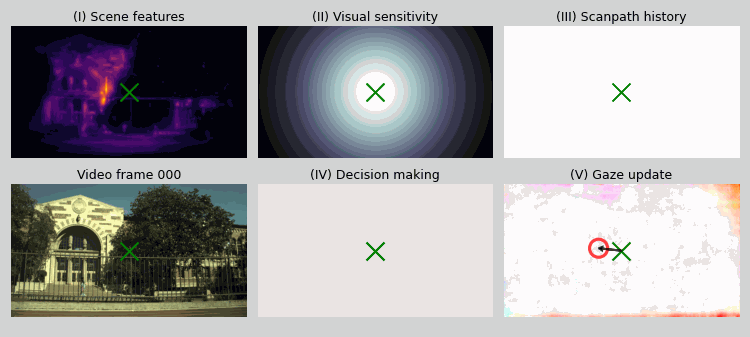

### S2_fig.tif

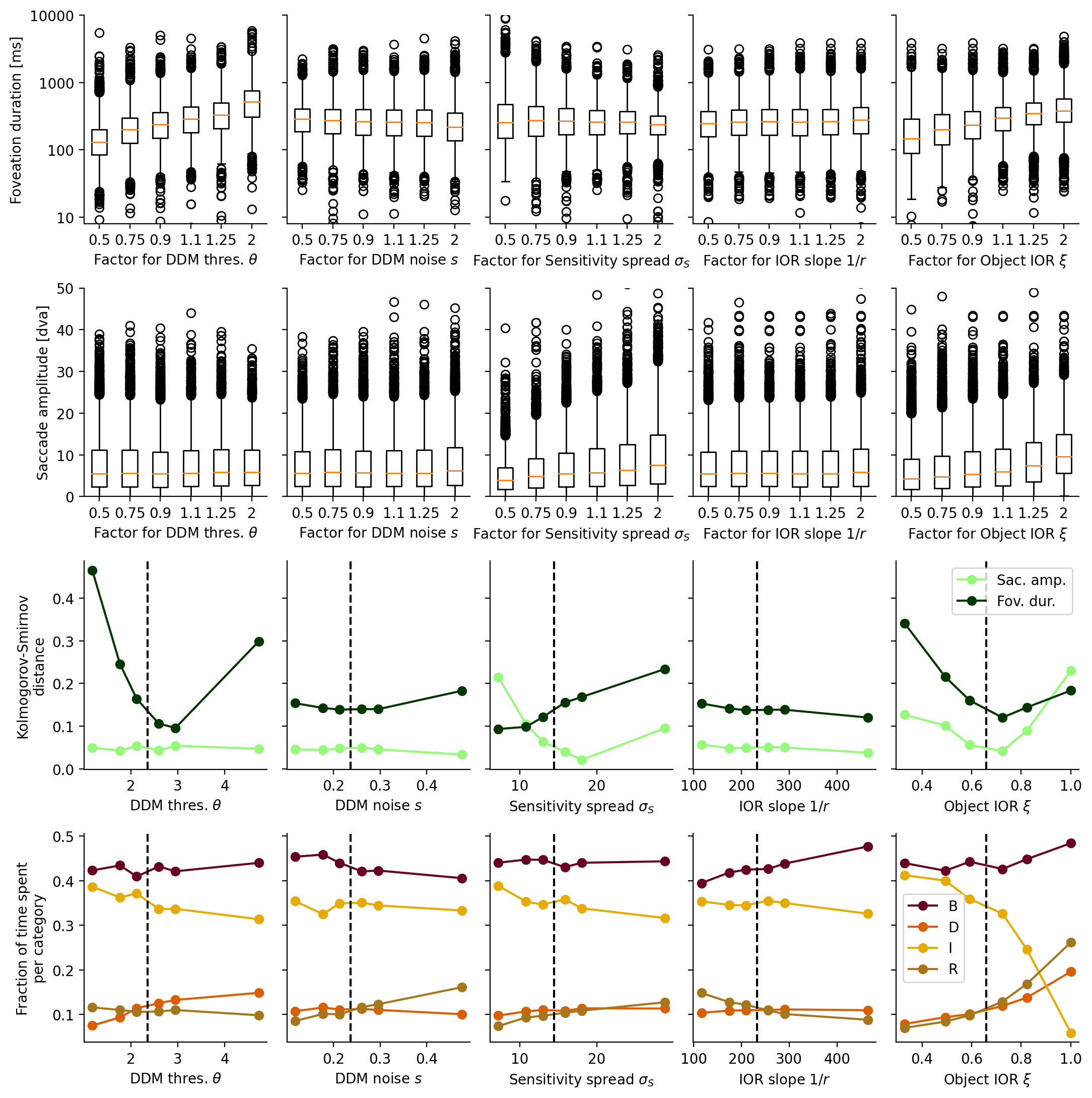

### S2_file.gif

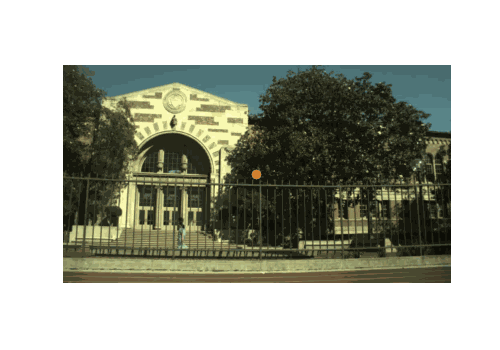

### S3_fig.tif

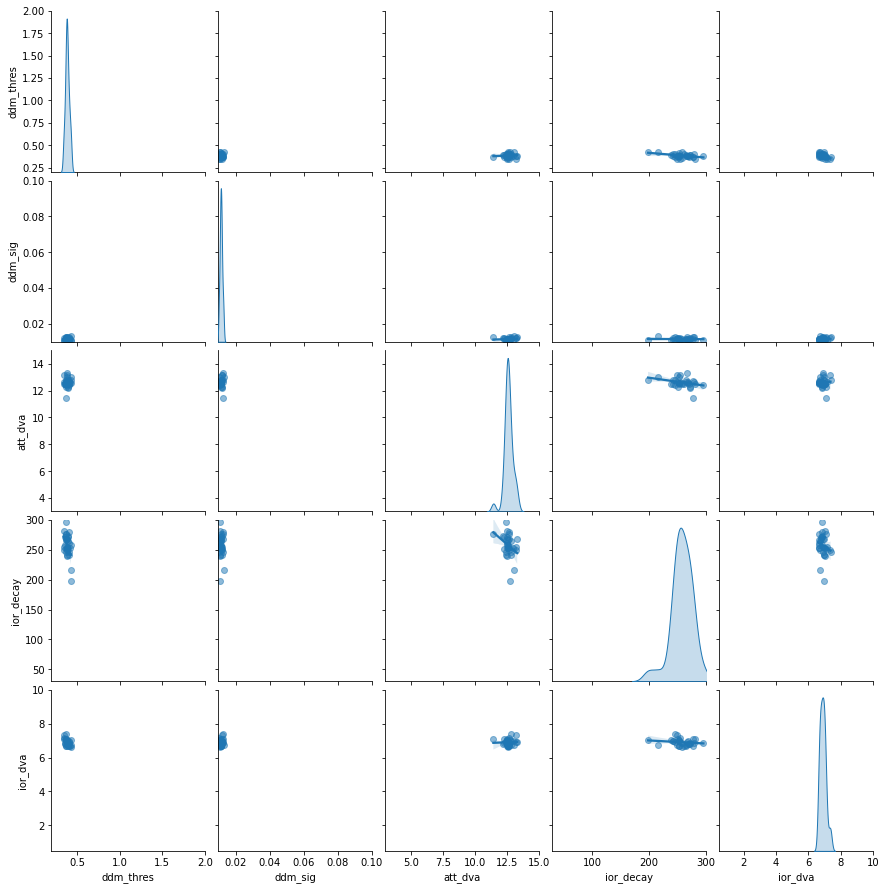

### S3_file.gif

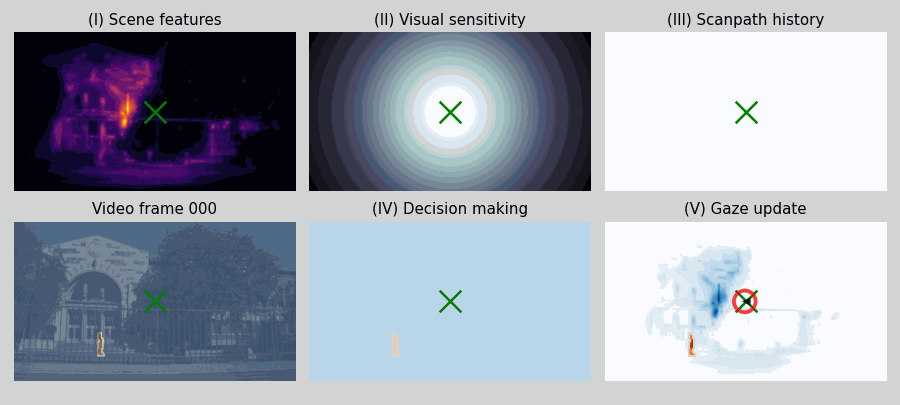

### S4_fig.tif

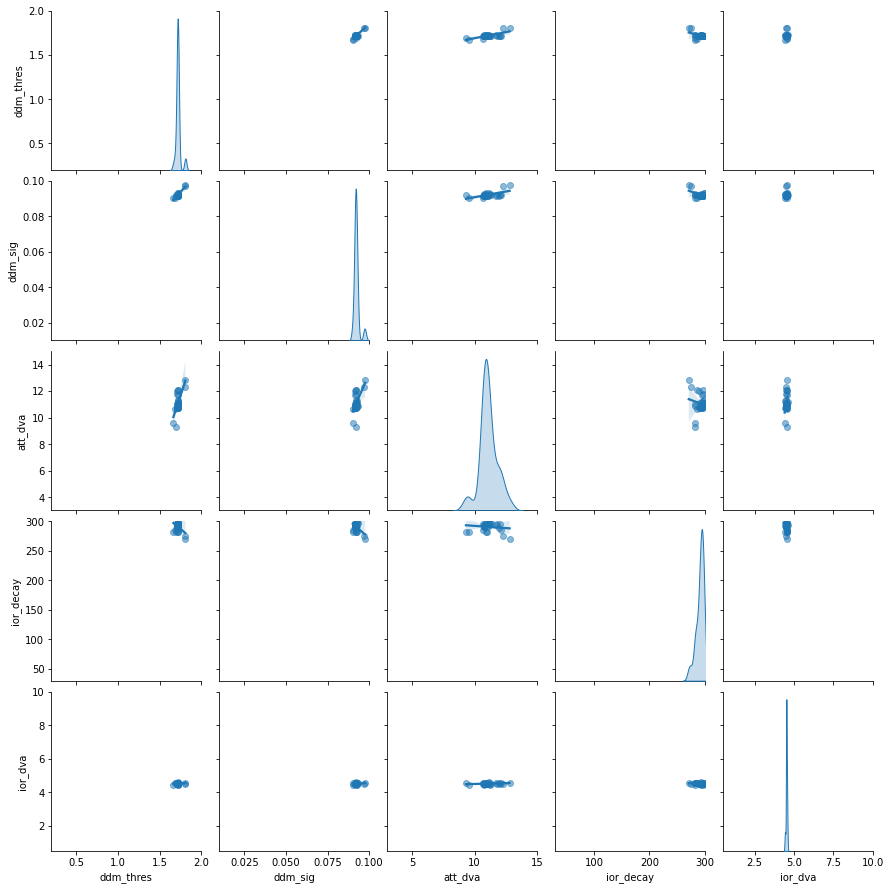

### S4_file.gif

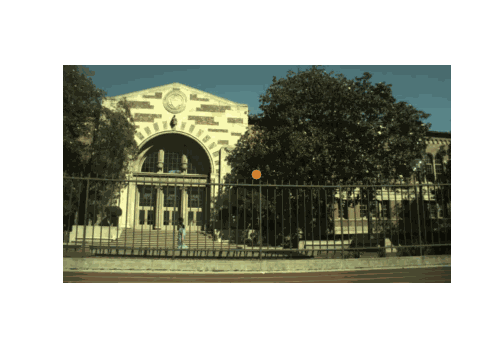

### S5_fig.tif

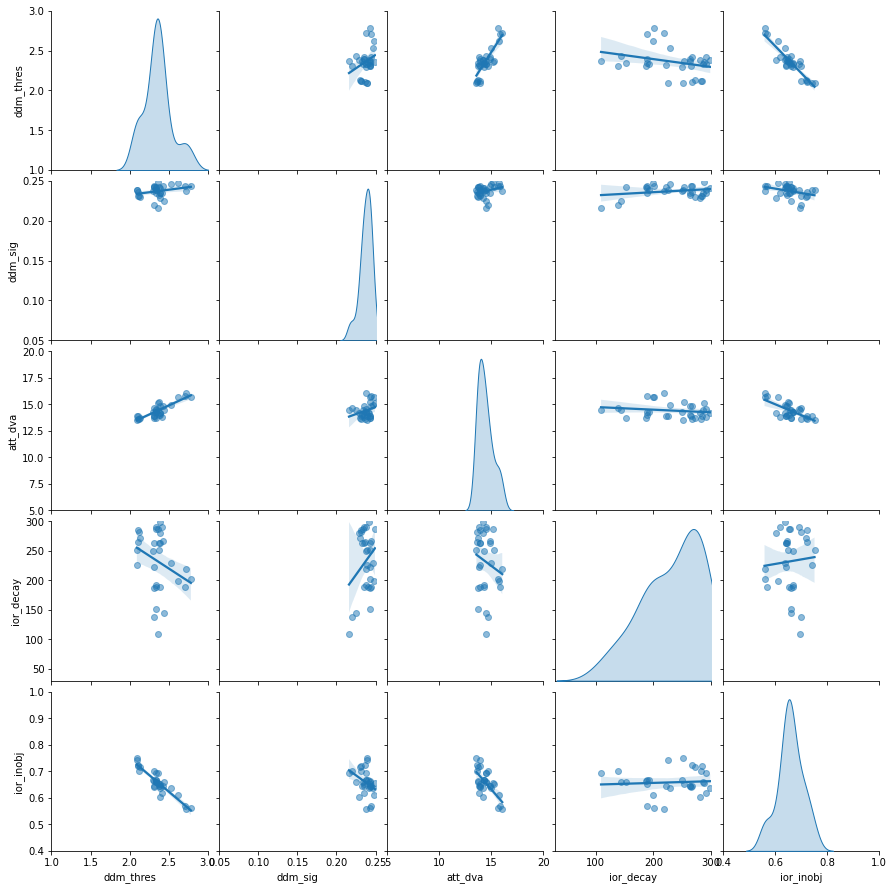

### S6_fig.tif

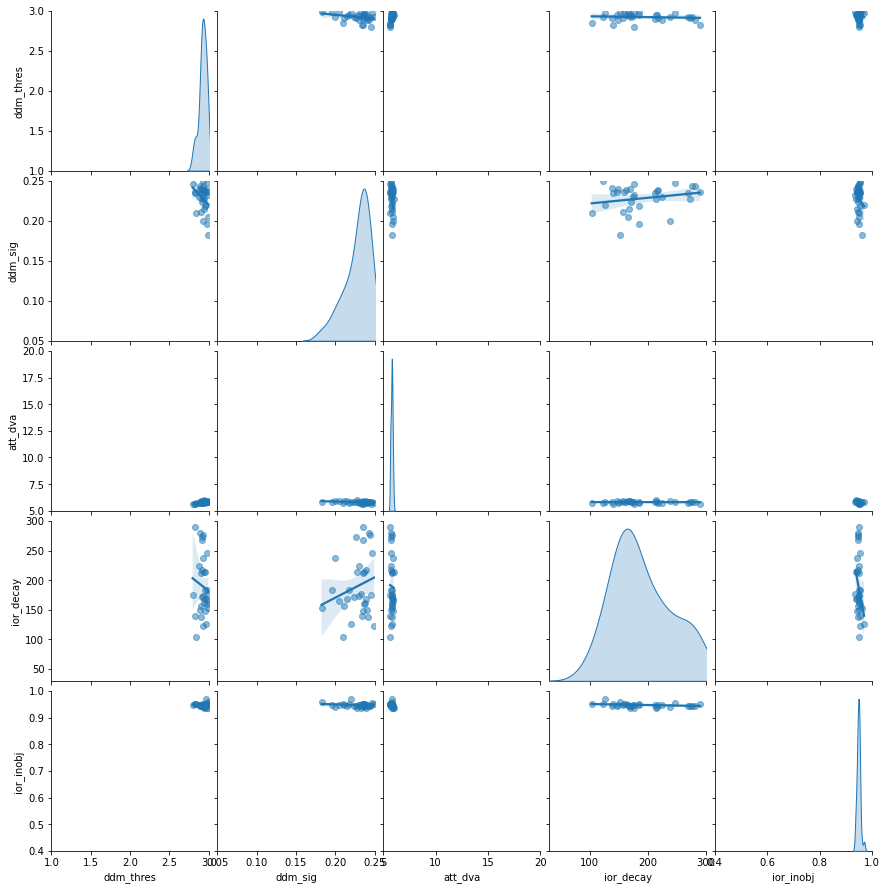

### S7_fig.tif

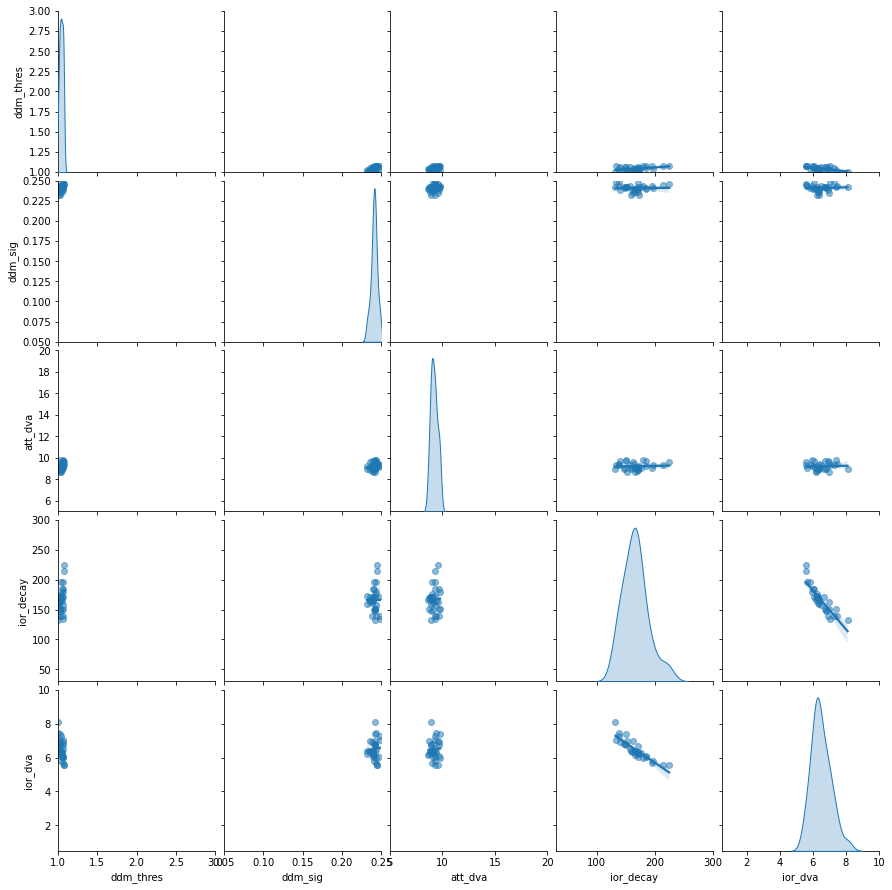

### S9_fig.tif

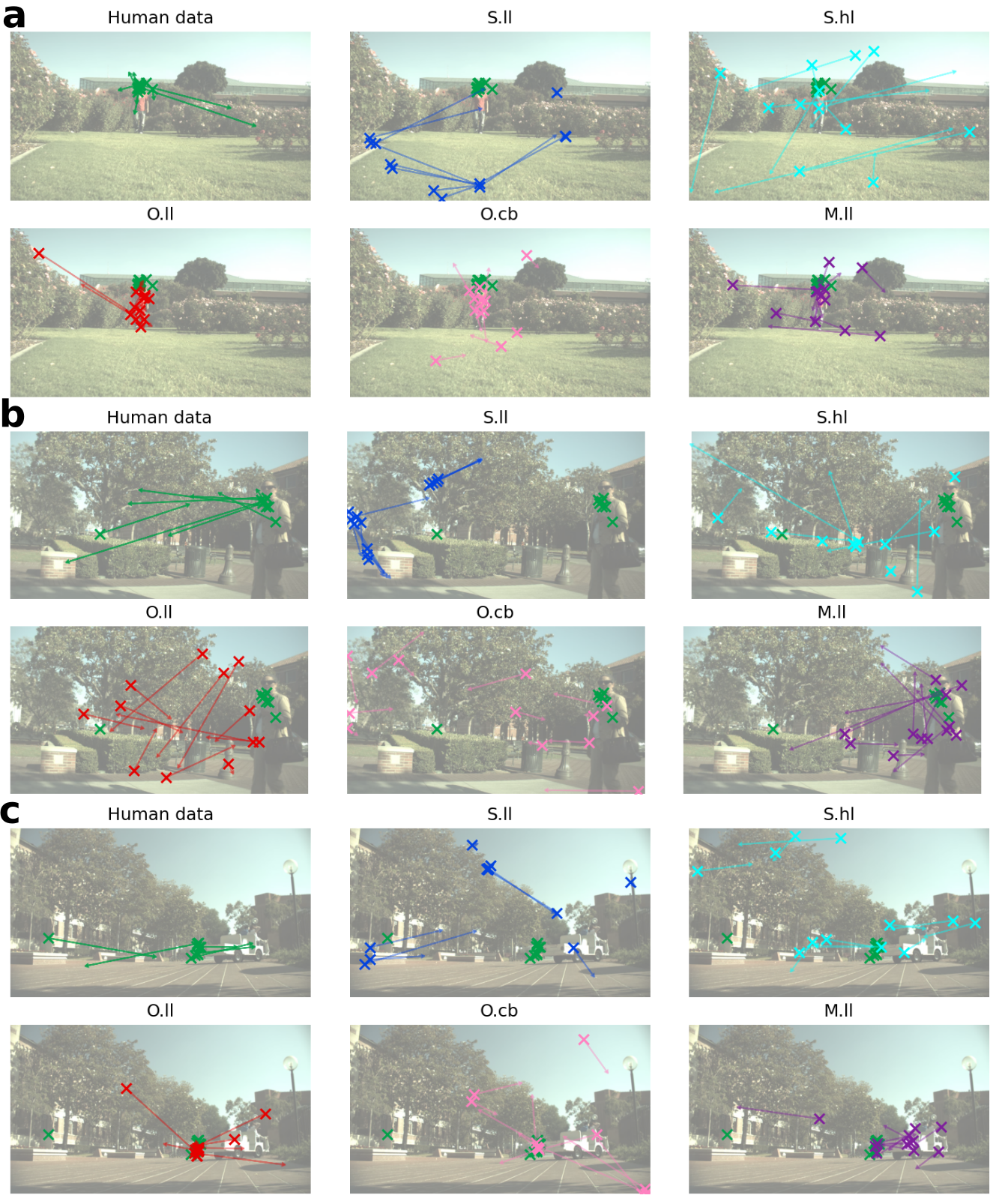
